# Highly Divergent Neuropeptide - non-coding RNA regulatory networks underpin variant host-finding behaviours in *Steinernema* species infective juveniles

**DOI:** 10.1101/2020.10.30.359141

**Authors:** Neil D. Warnock, Erwan Atcheson, Ciaran McCoy, Johnathan J. Dalzell

## Abstract

We conducted a transcriptomic and small RNA analysis of infective juveniles (IJs) from three behaviourally distinct *Steinernema* species. Substantial variation was found in the expression of shared gene orthologues, revealing gene expression signatures that correlate with behavioural states. 97% of predicted microRNAs are novel to each species. Surprisingly, our data provide evidence that isoform variation can effectively convert protein-coding neuropeptide genes into non-coding transcripts, which may represent a new family of long non-coding RNAs. These data suggest that differences in neuropeptide gene expression, isoform variation, and small RNA interactions could contribute to behavioural differences within the *Steinernema* genus.

---

*Steinernema* spp. nematodes are obligate entomopathogens that invade and kill insect hosts through coordinated action with commensal *Xenorhabdus* bacteria (Lu et al., 2017). *Steinernema* spp. infective juveniles (IJs) are developmentally arrested, non-feeding, and adapted for host-finding. The IJs display qualitatively different host-finding strategies between species (Spence et al., 2008). However we have also reported substantial behavioural variation between populations of *Steinernema carpocapsae* (Warnock et al., 2019). *S. carpocapsae* is classed as an ‘ambusher’, and will nictate and jump directionally in response to host-specific volatiles and mechanosensory input. *Steinernema glaseri* is classed as a ‘cruiser’, and does not nictate or jump, relying entirely on sinusoidal motility for host-finding and invasion. *Steinernema feltiae* employs a qualitatively intermediate strategy relative to the other two species. Whilst these behavioural categories represent a simplification of what manifests as a behavioural continuum within and between species, they provide a useful framework for the study of the molecular basis of behaviour, perhaps uniquely so for a single life-stage, within a single genus of the phylum Nematoda. The recent development of genomic and transcriptomic resources for a range of *Steinernema* spp. highlights efforts to develop these species as models for comparative biology (Dillman et al., 2015; Macchietto et al., 2017; Warnock et al., 2019).

In recent work, we demonstrated that behavioural variation between populations of *S. carpocapsae* correlated with variation in the expression of genes and microRNAs. In particular, a range of neuronal genes were implicated, including neuropeptides (J. S. Lee et al., 2017; Morris et al., 2017; Warnock et al., 2019). Neuropeptide gene expression is enriched within the IJ stage of numerous parasitic nematode species (J. S. Lee et al., 2017), suggesting a critical role in behavioural diversification, which is supported by functional investigation in *S. carpocapsae* (Morris et al., 2017).

For RNAseq analysis of three species – *Steinernema carpocapsae,* (strain ALL), *Steinernmea glaseri* (strain NC), and *Steinernema feltiae* (strain SN) maintained in *Galleria mellonella* – three biological replicates of each strain were prepared from approximately 10,000 individuals. The majority (80%) of neuronal genes detected were orthologous between all three species (Fig 1, Supplemental files 1-2). Considerable variation was seen in the expression levels of neuronal genes shared between species. To correlate with host-seeking behaviour, where *S. carpocapsae* is considered an ‘ambusher’, *S. glaseri* a ‘cruiser’ and *S. feltiae* intermediate in behaviour, differential expression between species would be expected to conform to one of two patterns: Sc>Sf>Sg or Sc<Sf<Sg. Expression levels of ten such peptides correlated with behaviour, with upregulation of *flp-11, flp-25, flp-33, nlp-14b, nlp-37, nlp-55* and *RYamide* positively correlated, and *nlp-3, nlp-36* and *snet-1* negatively correlated with ‘ambushing’ behaviour. Only one of these, *nlp-36*, showed a high level of differential expression (>2 log2-fold). Interestingly, *nlp-36* was previously found to negatively correlate with nictation behaviour between three *S. carpocapsae* strains (Warnock et al., 2019).

**Fig 1.**
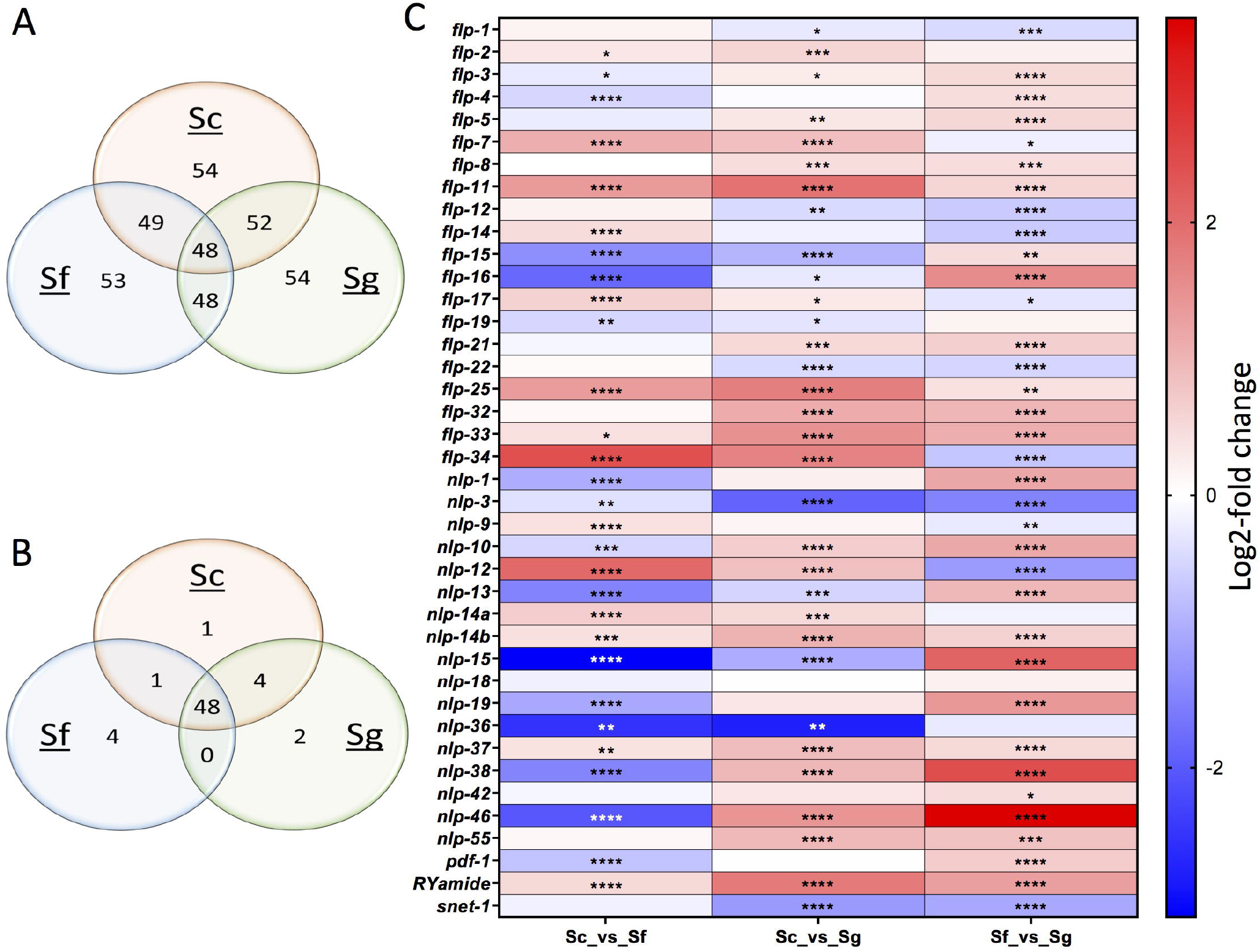
Venn diagrams showing (A) total or (B) number of *nlp* and *flp* genes unique to species, with (C) heatmap showing pairwise comparison of differentially expressed genes common across the three species. Three transcriptome libraries were prepared for each species (TruSeq RNA Library Prep Kit v2, Illumina). Sequencing was performed on the HiSeq2500 instrument. Data quality was assessed using FastQC, trimmed using Trimmomatic (Bolger et al., 2014), and mapped to the *S. carpocapsae* genome (Dillman et al., 2015) using the STAR (v. 2.5.3a) package (Dobin et al., 2013). Differential expression of genes was quantified using the DESeq2 (v. 1.14.1) package (Love et al., 2014). *Steinernema* spp. neuropeptide gene homologues were identified via reciprocal BLAST analysis. The top reciprocal BLAST hit from *C. elegans* was used to assign putative neuropeptide gene names, and subsequent manual comparison of *Steinernema* neuropeptide primary sequences to established nematode neuropeptide motifs (McVeigh et al., 2008; 2005) was used to confirm or reassign gene names where appropriate. ****, p<0.0001, *** p<0.001, ** p<0.01, *p<0.05.

Small RNA libraries were generated from infective juveniles of the three *Steinernema* species. MicroRNAs are small non-coding RNAs that regulate the expression of target genes. Considerable research effort has focused on the role of microRNAs in development, but increasingly, studies are linking microRNAs and other small RNAs to diverse aspects of biology, including behaviour (Cox et al., 2019; Warnock et al., 2019). A total of 930 microRNAs were identified across all three species. Strikingly, a very small proportion (31/930, 3%) of microRNAs were identical in all three species (Fig 2, Supplemental files 1 and 3). Each *Steinernema* species showed a distinct expression profile for these microRNAs. Seven microRNAs correlated with behaviour, with *miR-19-3p, miR-3-3p,* and *miR-410-5p* positively, and *miR-352-5p, miR-368-5p, miR-417-3p* and *miR-767-3p* negatively associated with ‘ambusher’ behaviour. Predicted microRNA binding sites were identified using miRanda (Enright et al., 2003). Two separate miRanda analyses were performed, using i) unrestricted settings and ii) strict settings that require perfect conservation of seed site sequence complementarity between microRNA and target mRNA (Supplemental files 3-6). *miR-19-3p* and *miR-410-5p* are predicted to target *S. glaseri nlp-36*; consistent with the association of *nlp-36* expression with ‘cruiser’ behaviour, these two miRNAs are upregulated in ‘ambushers’, which should have the effect of further suppressing *nlp-36* expression, lending further support to the hypothesis that this neuropeptide is functionally involved in *Steinernema* host-seeking behaviour. Similarly, *miR-352-5p*, downregulated in ‘ambushers’, targets *S. feltiae flp-25*, upregulated in ‘ambushers’. Finally, *miR-417-3p* and *miR-767-3p*, both upregulated in ‘cruisers’, target *S. carpocapsae flp-11*, downregulated in ‘cruisers’. Together the miRNA evidence lends support to *nlp-36, flp-25*, and *flp-11* expression levels being important determinants of host-seeking behaviour.

**Fig 2.**
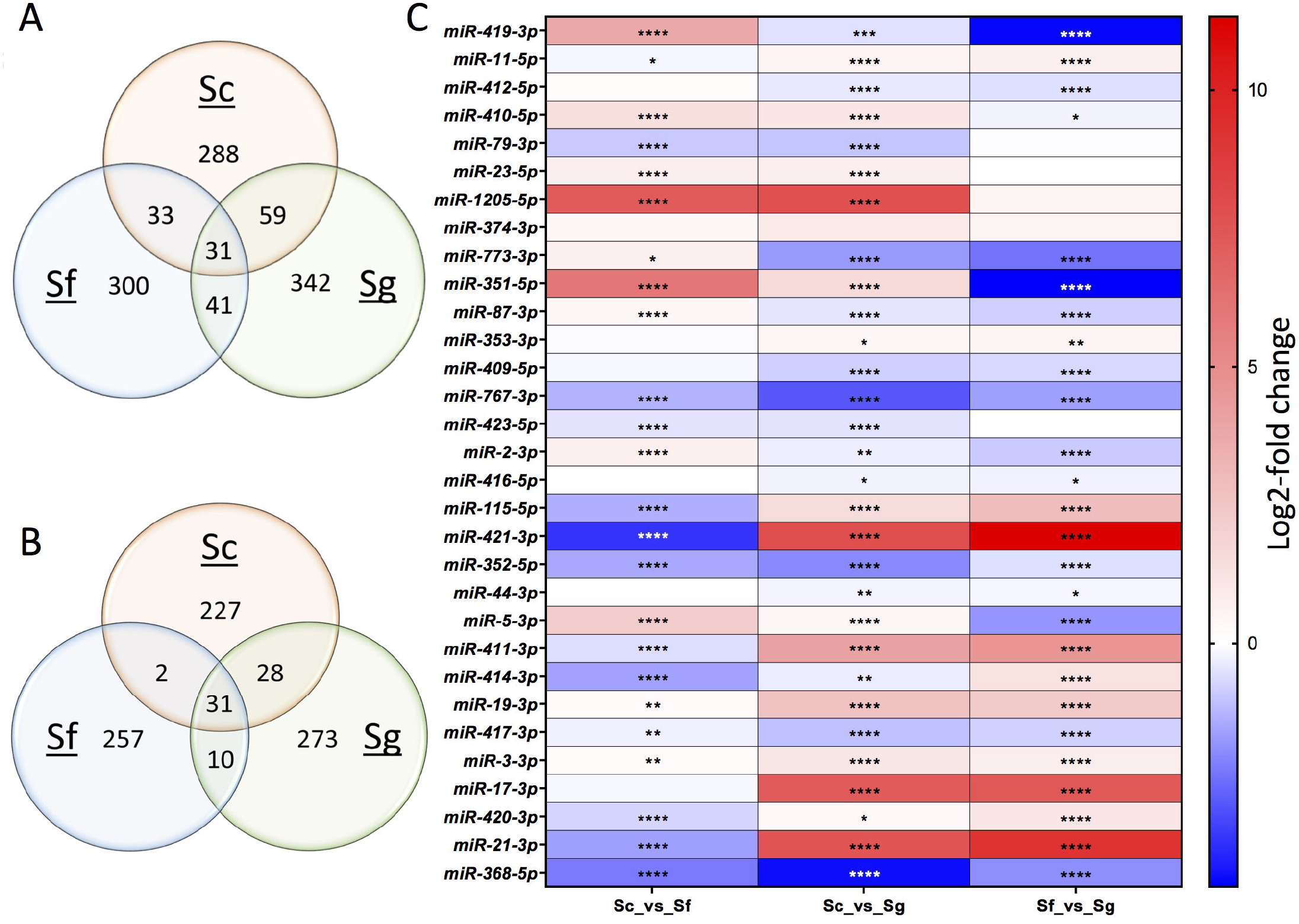
Venn diagrams showing (A) total or (B) number of microRNAs unique to species, with (C) heatmap showing pairwise comparison of differentially expressed microRNAs common across the three species. Three small RNA libraries were prepared for each species using the TruSeq Small RNA library Kit (Illumina), and 50 bp single-end libraries were sequenced on the HiSeq2500 instrument. Quality assessment was performed using FastQC, trimming by Cutadapt. Reads that passed QC were mapped to the genome sequence of *S. carpocapsae*, and microRNAs were identified by miRDeep2 (v. 2.0.0.8) (Friedländer et al., 2012), using a training set of mature and precursor microRNA sequences downloaded from miRBase. Naming of microRNAs was preferentially aligned with *C. elegans*, as indicated by miRDeep2 output. Differentially expressed microRNAs were identified using the DESeq2 package. ****, p<0.0001, *** p<0.001, ** p<0.01, *p<0.05.

As neuropeptides have been linked to behavioural variation in nematode infective juveniles, we assessed the degree to which the complexity of neuropeptide gene isoform complements might underpin behavioural strategy, using PacBio to sequence full length transcripts. SMRTbell library preparation and long read Isoform sequencing (PacBio Iso-seq) was performed at the Earlham Institute (Norwich, UK). This brought to light the complexity of isoforms for neuropeptide genes in the three *Steinernema* species. Isoform variation increased in line with the complexity of host-finding behaviour in these species, with 72, 62 and 59 such isoforms in *S. carpocapsae, S. feltiae* and *S. glaseri* respectively (Supplemental file 7). Furthermore, for many neuropeptide genes it was discovered that long non-coding isoforms of the functional transcripts exist (Fig 3, Supplemental file 7). Altogether one third (66 of 193) of the identified neuropeptide transcripts were non-coding. This could represent another level of gene regulation. For example, the *flp-11* non-coding transcript is the predicted target of *miR-417-3p* and *miR-767-3p* microRNAs. *S. carpocapsae flp-11* is downregulated in ‘cruisers’, and associated with ‘ambushing’. Not only are *miR-417-3p* and *miR-767-3p* downregulated in *S. carpocapsae*, but their capacity to reduce *flp-11* expression appears to be further diluted by the presence of an *flp-11* isoform in *S. carpocapsae* which is non-coding and which is presumably able to compete with the functional *flp-11* transcript for binding of *miR-417-3p* and *miR-767-3p.* A similar pattern of expression was observed with *flp-25* isoforms.

**Fig 3.**
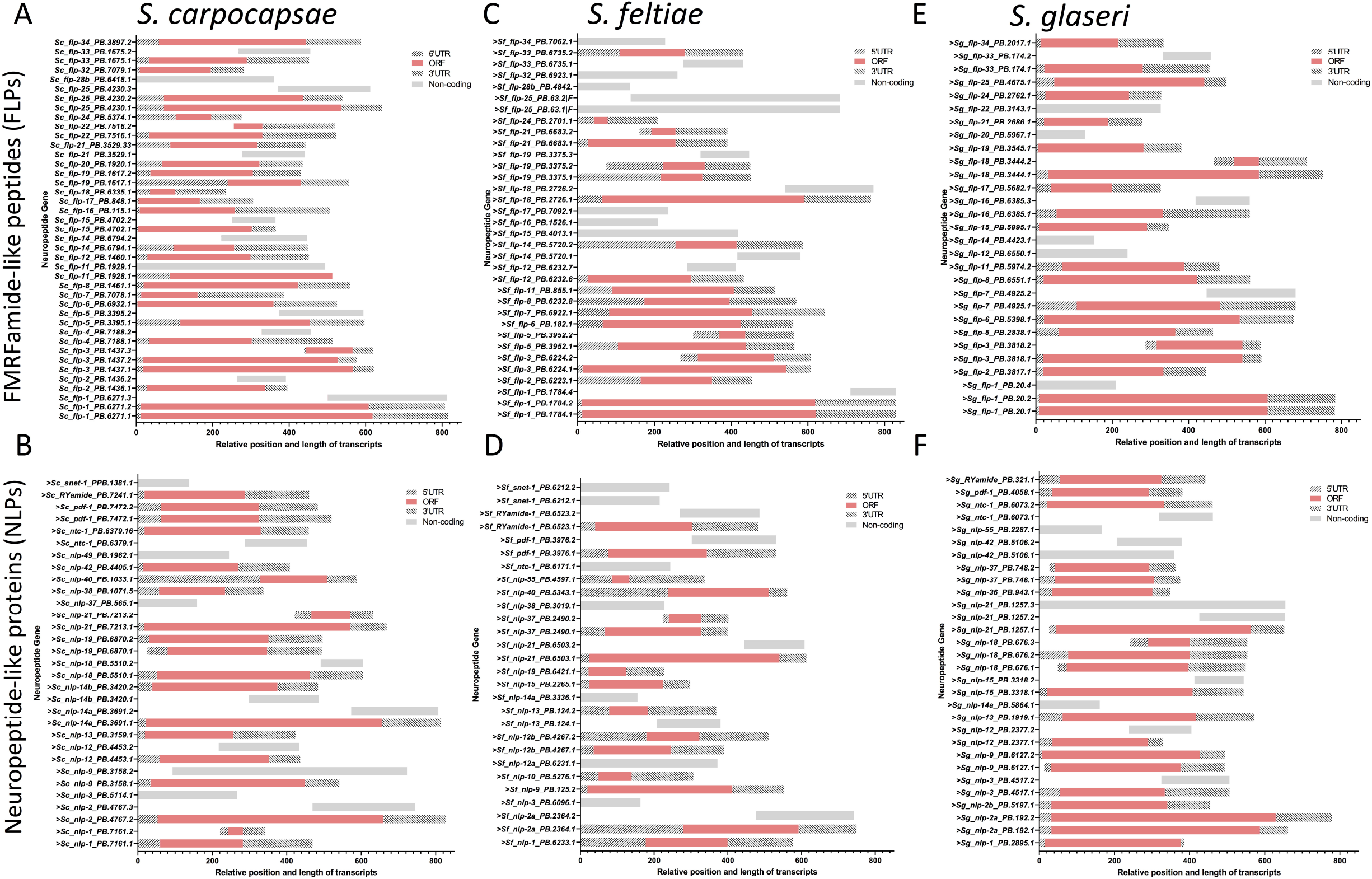
Schematic representation of *flp* and *nlp* neuropeptide gene transcripts, with either UTRs and ORFs, or as non-coding transcripts, for each isoform. (A) *flp* and (B) *nlp* isoforms from *S. carpocapsae*; (C) *flp* and (D) *nlp* isoforms from *S. feltiae; (F-)flp* and (F) *nlp* isoforms from *S. glaseri*. The Iso-Seq analysis pipeline was run via the command-line tools only option of the SMRT link software installation. Raw subreads were converted into error corrected Cluster Consensus Sequences and then classified as full-length or non-full-length reads, artificial-concatemer chimeric, or non-chimeric reads. Iterative Clustering and Error correction (ICE) and Quiver were used to further group and polish these sequences, and aligned to the genome using STAR alignment software. Cupcake Tofu associated python scripts were employed to collapse any redundant isoforms and count the number of CCSs that support each unique isoform (Gordon et al., 2015). Positioning is relative to beginning of alignment with longest isoform.

In summary, these data suggest that a complex system of neuropeptide gene regulation may underly behavioural differences across three species of *Steinernema*. The expression profiles of several neuropeptide genes correlate with behavioural differences, as does expression of microRNAs that are predicted to target those transcripts. The vast majority (97%) of microRNAs are unique to each species. Another layer of regulation is potentially mediated by the surprising ubiquity of long non-coding isoforms of almost all expressed neuropeptide genes within each species, opening the possibility that competition of microRNA binding to these non-coding transcripts also affects transcript levels of key neuropeptide genes involved in behaviour, and contributing to the differences in host-seeking behaviour seen in these three species.

Expression levels of ten neuropeptides genes correlated with behaviour across the three species, of which three, *nlp-36, flp-11* and *flp-25* showed the best evidence across the study of playing a potentially functional role. *nlp-36* was previously shown to correlate with decreased nictation behaviour across three *S. carpocapsae* strains (Warnock et al., 2019); the current finding that expression of this gene also correlates with decreased nictation behaviour across three species of *Steinernema* reinforces the evidence that *nlp-36* is crucial to these differences. In the model nematode *Caenorhabditis elegans* there is evidence that a cyclic nucleotide-gated channel subunit, TAX-2, which positively regulates NLP-36, is involved in thermosensation, olfaction, chemosensation, and axon guidance (Coburn and Bargmann, 1996; Coburn et al., 1998); differences in host-seeking behaviour could be mediated via *nlp-36* expression by any of these pathways. Curiously, the FLP-11 neuropeptide, functionally associated in *C. elegans* with the systemic onset of sleep and locomotor cessation (Steuer Costa et al., 2019; Turek et al., 2016), in *Steinernema* is associated with ‘ambushing’ host-seeking behaviour; although in a different nematode species, *Ascaris suum*, FLP-11 induces a quite different response, increasing contraction frequency of the ovijector (Moffett et al., 2003). Besides decreased dauer entry in an *flp-25* mutant of *C. elegans* (J. S. Lee et al., 2017), there appear to be few functional studies of FLP-25.

The role of microRNAs in regulating gene expression and attendant effects on behaviour is well-supported, and the complexity of these systems increasingly appreciated (Gillan et al., 2017; Gu et al., 2017; D. Lee et al., 2017). This study has shown considerable variation in microRNA expression between species, and identified several microRNAs that correlate with host-finding behaviour in *Steinernema*. Furthermore, several of the predicted target genes of these microRNAs were also identified as correlating with host-finding behaviour, in ways consistent with their proposed functional roles. However, these associations remain hypothetical. Further work will be required to validate the specificity of microRNAs with their predicted transcript targets, for instance by co-localisation of microRNAs and transcripts, and via Argonaute CLIP-seq (Zisoulis et al., 2010). In parallel, confirming the functionality of genes and microRNAs here identified as associated with behaviour will require genetic and molecular manipulation such as knock-down or knockout of those particular genes and transcripts.

A previous study of the genetic basis of behavioural difference in *S. carpocapsae* strains found evidence of an interaction between isoform variation and attendant microRNA targeting influencing behaviour (Warnock et al., 2019). Here for the first time long non-coding isoforms of neuropeptide transcripts are shown to be the targets of microRNAs associated with different host-finding behaviours in three *Steinernema* species. To our knowledge, this is the first such discovery in any organism. If validated it could represent a novel additional mechanism of microRNA regulation of neuropeptide gene transcripts. Particularly compelling is the presence in *S. carpocapsae* of a non-coding isoform of *flp-11*, shown to associate with ‘ambushing’ behaviour. This isoform is predicted to be the target of two microRNAs themselves associated with the ‘cruiser’ form of host-seeking behaviour, and hence the non-coding isoform could be competing for these microRNAs further reducing their effect on *flp-11* transcript levels, which ought then to result in increased translation of these transcripts. If this gene genuinely plays a functional role in the ‘ambusher’ behaviour in this species, the non-coding isoform could be contributing to this behaviour by the mechanism outlined.

This is the first study to associate differentially-expressed genes correlating with host-seeking behaviours in three *Steinernema* species. The complexity of gene regulation is revealed by the number of microRNAs uniquely or differentially expressed by each species. In addition, a third level of gene regulation is unexpectedly revealed in the form of long non-coding isoforms of neuropeptide genes, predicted to compete for binding of microRNAs also associated with different forms of host-seeking behaviour.

## Acknowledgements

Waxworms infected with *S.feltiae* (SN), and *S. glaseri* (NC) were kindly provided by Ali Mortazavi and Marissa Macchietto, University of California, Irvine.

The authors declare no competing interests.

## Supporting information captions

**Supplemental file 1.** Comparison of all differentially expressed genes and microRNAs shared between all three *Steinernema* species.

**Supplemental file 2.** Counts of all differentially expressed genes shared between three *Steinernema* species.

**Supplemental file 3.** Counts of all differentially expressed microRNAs shared between three *Steinernema* species.

**Supplemental file 4.** Miranda output file for microRNA targets in *S. carpocapsae*. **Supplemental file 5.** Miranda output file for microRNA targets in *S. feltiae*.

**Supplemental file 6.** Miranda output file for microRNA targets in *S. glaseri*.

**Supplemental file 7.** Table of neuropeptide transcript isoforms from PacBio sequencing

Transcriptomic and small RNA raw data, and circular consensus sequences from PacBio sequencing, available at NCBI accession number PRJNA662954

## References

Bolger, A.M., Lohse, M., Usadel, B., 2014. Trimmomatic: a flexible trimmer for Illumina sequence data. Bioinformatics 30, 2114–2120. doi:10.1093/bioinformatics/btu170

Coburn, C.M., Bargmann, C.I., 1996. A putative cyclic nucleotide-gated channel is required for sensory development and function in C. elegans. Neuron 17, 695–706. doi:10.1016/s0896-6273(00)80201-9

Coburn, C.M., Mori, I., Ohshima, Y., Bargmann, C.I., 1998. A cyclic nucleotide-gated channel inhibits sensory axon outgrowth in larval and adult Caenorhabditis elegans: a distinct pathway for maintenance of sensory axon structure. Development 125, 249–258.

Cox, D., Reilly, B., Warnock, N.D., Dyer, S., Sturrock, M., Cortada, L., Coyne, D., Maule, A.G., Dalzell, J.J., 2019. Transcriptional signatures of invasiveness in Meloidogyne incognita populations from sub-Saharan Africa. Int. J. Parasitol. 49, 837–841. doi:10.1016/j.ijpara.2019.05.013

Dillman, A.R., Macchietto, M., Porter, C.F., Rogers, A., Williams, B., Antoshechkin, I., Lee, M.- M., Goodwin, Z., Lu, X., Lewis, E.E., Goodrich-Blair, H., Stock, S.P., Adams, B.J., Sternberg, P.W., Mortazavi, A., 2015. Comparative genomics of Steinernema reveals deeply conserved gene regulatory networks. Genome Biol. 16, 200–21. doi: 10.1186/s13059-015-0746-6

Dobin, A., Davis, C.A., Schlesinger, F., Drenkow, J., Zaleski, C., Jha, S., Batut, P., Chaisson, M., Gingeras, T.R., 2013. STAR: ultrafast universal RNA-seq aligner. Bioinformatics 29, 15–21. doi:10.1093/bioinformatics/bts635

Enright, A.J., John, B., Gaul, U., Tuschl, T., Sander, C., Marks, D.S., 2003. MicroRNA targets in Drosophila. Genome Biol. 5, R1–14. doi:10.1186/gb-2003-5-1-r1

Friedländer, M.R., Mackowiak, S.D., Li, N., Chen, W., Rajewsky, N., 2012. miRDeep2 accurately identifies known and hundreds of novel microRNA genes in seven animal clades. Nucleic Acids Res. 40, 37–52. doi:10.1093/nar/gkr688

Gillan, V., Maitland, K., Laing, R., Gu, H., Marks, N.D., Winter, A.D., Bartley, D., Morrison, A., Skuce, P.J., Rezansoff, A.M., Gilleard, J.S., Martinelli, A., Britton, C., Devaney, E., 2017. Increased Expression of a MicroRNA Correlates with Anthelmintic Resistance in Parasitic Nematodes. Front Cell Infect Microbiol 7, 452. doi:10.3389/fcimb.2017.00452

Gordon, S.P., Tseng, E., Salamov, A., Zhang, J., Meng, X., Zhao, Z., Kang, D., Underwood, J., Grigoriev, I.V., Figueroa, M., Schilling, J.S., Chen, F., Wang, Z., 2015. Widespread Polycistronic Transcripts in Fungi Revealed by Single-Molecule mRNA Sequencing. PLoS ONE 10, e0132628. doi:10.1371/journal.pone.0132628

Gu, H.Y., Marks, N.D., Winter, A.D., Weir, W., Tzelos, T., McNeilly, T.N., Britton, C., Devaney, E., 2017. Conservation of a microRNA cluster in parasitic nematodes and profiling of miRNAs in excretory-secretory products and microvesicles of Haemonchus contortus. PLoS Negl Trop Dis 11, e0006056. doi:10.1371/journal.pntd.0006056

Lee, D., Yang, H., Kim, J., Brady, S., Zdraljevic, S., Zamanian, M., Kim, H., Paik, Y.-K., Kruglyak, L., Andersen, E.C., Lee, J., 2017. The genetic basis of natural variation in a phoretic behavior. Nat Commun 8, 273–7. doi:10.1038/s41467-017-00386-x

Lee, J.S., Shih, P.-Y., Schaedel, O.N., Quintero-Cadena, P., Rogers, A.K., Sternberg, P.W., 2017. FMRFamide-like peptides expand the behavioral repertoire of a densely connected nervous system. Proc. Natl. Acad. Sci. U.S.A. 114, E10726–E10735. doi:10.1073/pnas.1710374114

Love, M.I., Huber, W., Anders, S., 2014. Moderated estimation of fold change and dispersion for RNA-seq data with DESeq2. Genome Biol. 15, 550–21. doi:10.1186/s13059-014-0550-8

Lu, D., Macchietto, M., Chang, D., Barros, M.M., Baldwin, J., Mortazavi, A., Dillman, A.R., 2017. Activated entomopathogenic nematode infective juveniles release lethal venom proteins. PLoS Pathog. 13, e1006302. doi:10.1371/journal.ppat.1006302

Macchietto, M., Angdembey, D., Heidarpour, N., Serra, L., Rodriguez, B., El-Ali, N., Mortazavi, A., 2017. Comparative Transcriptomics of Steinernema and Caenorhabditis Single Embryos Reveals Orthologous Gene Expression Convergence during Late Embryogenesis. Genome Biol Evol 9, 2681–2696. doi:10.1093/gbe/evx195

McVeigh, P., Alexander-Bowman, S., Veal, E., Mousley, A., Marks, N.J., Maule, A.G., 2008. Neuropeptide-like protein diversity in phylum Nematoda. Int. J. Parasitol. 38, 1493–1503. doi:10.1016/j.ijpara.2008.05.006

McVeigh, P., Kimber, M.J., Novozhilova, E., Day, T.A., 2005. Neuropeptide signalling systems in flatworms. Parasitology 131 Suppl, S41–55. doi:10.1017/S003H82005008851

Moffett, C.L., Beckett, A.M., Mousley, A., Geary, T.G., Marks, N.J., Halton, D.W., Thompson, D.P., Maule, A.G., 2003. The ovijector of Ascaris suum: multiple response types revealed by Caenorhabditis elegans FMRFamide-related peptides. Int. J. Parasitol. 33, 859–876. doi: 10.1016/s0020-7519(03)00109-7

Morris, R., Wilson, L., Sturrock, M., Warnock, N.D., Carrizo, D., Cox, D., Maule, A.G., Dalzell, J.J., 2017. A neuropeptide modulates sensory perception in the entomopathogenic nematode Steinernema carpocapsae. PLoS Pathog. 13, e1006185. doi:10.1371/journal.ppat.1006185

Spence, K.O., Lewis, E.E., Perry, R.N., 2008. Host-finding and invasion by entomopathogenic and plant-parasitic nematodes: evaluating the ability of laboratory bioassays to predict field results. J. Nematol. 40, 93–98.

Steuer Costa, W., Van der Auwera, P., Glock, C., Liewald, J.F., Bach, M., Schüler, C., Wabnig, S., Oranth, A., Masurat, F., Bringmann, H., Schoofs, L., Stelzer, E.H.K., Fischer, S.C., Gottschalk, A., 2019. A GABAergic and peptidergic sleep neuron as a locomotion stop neuron with compartmentalized Ca2+ dynamics. Nat Commun 10, 4095–15. doi: 10.1038/s41467-019-12098-5

Turek, M., Besseling, J., Spies, J.-P., König, S., Bringmann, H., 2016. Sleep-active neuron specification and sleep induction require FLP-11 neuropeptides to systemically induce sleep. Elife 5, 287. doi:10.7554/eLife.12499

Warnock, N.D., Cox, D., McCoy, C., Morris, R., Dalzell, J.J., 2019. Transcriptional variation and divergence of host-finding behaviour in Steinernema carpocapsae infective juveniles. BMC Genomics 20, 884–17. doi:10.1186/s12864-019-6179-y

Zisoulis, D.G., Lovci, M.T., Wilbert, M.L., Hutt, K.R., Liang, T.Y., Pasquinelli, A.E., Yeo, G.W., 2010. Comprehensive discovery of endogenous Argonaute binding sites in Caenorhabditis elegans. Nat. Struct. Mol. Biol. 17, 173–179. doi:10.1038/nsmb.1745

